# Loss of one Engrailed1 allele enhances induced α-synucleinopathy

**DOI:** 10.1101/530915

**Authors:** Diptaman Chatterjee, Daniel Saiz Sanchez, Emmanuel Quansah, Nolwen L Rey, Sonia George, Katelyn Becker, Zachary Madaj, Jennifer A Steiner, Jiyan Ma, Martha L Escobar Galvis, Jeffrey H Kordower, Patrik Brundin

## Abstract

**Background:** Parkinson’s disease (PD) is a synucleinopathy that has multiple neuropathological characteristics, with nigrostriatal dopamine system degeneration being a core feature. Current models of PD pathology typically fail to recapitulate several attributes of the pathogenic process and neuropathology. We aimed to define the effects of combining a mouse model exhibiting multiple PD-like changes with intrastriatal injections of α-synuclein (α-syn) pre-formed fibrils (PFFs) aggregates. We employed the heterozygous Engrailed 1 (*En1*^+/-^) mouse that features several pathophysiological hallmarks of clinical PD. Objective: To test the hypothesis that the neuropathological changes in the *En1*^+/-^ mice will promote formation of α-syn aggregates following intrastriatal injections of pathogenic human α-syn PFFs. Methods: We unilaterally injected PFFs into the striata of 1 month-old *En1*^+/-^ and control wild-type mice and euthanized animals at 3 months for post-mortem analysis. Results: Using immunohistochemistry and unbiased stereology, we established that PFF-injected *En1*^+/-^ mice exhibited a near-threefold increase in pS129-α-syn-positive neurons in the substantia nigra compared to PFF-injected wild-type mice. The PFF-injected *En1*^+/-^ mice also displayed significant increases in pS129-α-syn-positive neurons in the amygdala and ventral tegmental area; regions of known PD pathology with projections to the striatum. Additionally, we observed amplified pS129-α-syn-positive aggregation in *En1*^+/-^ mice in multiple cortical regions. Conclusions: Following intrastriatal injection of PFFs, absence of an *En1* allele leads to additional aggregation of pathological α-syn, potentially due to *En1*-loss mediated nigrostriatal impairment. We propose that further development of this double-hit model could result in a PD mouse model that predicts which experimental therapies will be effective in PD.

## Introduction

Parkinson’s disease (PD) is a synucleinopathy where degeneration of nigrostriatal dopamine neurons is a core neuropathological feature. It is believed that mitochondrial deficits and aggregation of misfolded α-synuclein (α-syn) are key components of the pathogenic process[1, 2]. Mitochondrial insults, particularly complex I deficiencies, have long been suggested to contribute to the development of PD pathology[3]. Toxins that cause complex I impairment, e.g. 1-methyl-4-phenyl-1,2,3,6-tetrahydropyridine (MPTP), have been used to generate animal models of PD, but these typically cause relatively acute degeneration of nigral dopaminergic neurons that is not representative of the progressive pathology of PD. α-Syn pathology has in a few cases been observed in MPTP-treated animals, but they are not a consistent feature of these models [4]. Furthermore, rotenone injections can lead to nigral neuron degeneration, as well as some α-syn pathology, but reproducibility can be problematic due to high rodent mortality [5, 6].

We [7] and others [8, 9] have previously characterized a PD-like neuropathology in the Engrailed1 heterozygous mouse model (*En1*^+/-^) including mitochondrial deficits, autophagic disturbances, axonal degeneration, neuroinflammation, diminished potassium-evoked dopamine release in the dorsal striatum and progressive loss of midbrain dopamine neurons. Engrailed1 has been shown to contribute to axonal maintenance and mitochondrial support in dopaminergic neurons into adulthood and counteract deficits associated with experimentally induced neuronal insults (MPTP, 6-OHDA, α-syn), both at the level of transcriptional regulation and translational output essential for mitochondrial function [8–13]. The primary aim of the present study was to define the effect of combining the complex neuropathology associated seen in *En1*^+/-^ mice with the progressive proteinopathy that follows intracerebral injection of α-syn fibrils [14]. Thus, we injected α-syn preformed fibrils (PFFs) into the striatum of *En1*^+/-^ mice. Eight weeks later we found significantly greater numbers of phosphorylated serine-129 α-syn (p-S129)-positive inclusions in the substantia nigra (SN) and ventral tegmental area (VTA), as well as in the basolateral amygdala (BLA) of male mice when compared to wild-type (WT) mice injected with the same amount of PFFs. We believe this novel *En1*-PFF mouse model represents a paradigm that mimics several aspects of cardinal PD pathologies. We believe that it can prove to be a highly relevant model system in which one can test experimental therapeutics aimed at slowing PD progression and obtain results that are highly predictive of clinical success.

## Methods and materials

### Animals

*En1*^+/-^ mice were housed 4-5 per cage under a regular cycle (12 h light/12 h dark) and had access to food and water *ad libitum.* The mice were treated in accordance with the Guide for the Care and Use of Laboratory Animals (National Institutes of Health) and all procedures and experimental conditions were undertaken with the prior approval of the Institutional Animal Use and Care Committee of Van Andel Research Institute. The *En1*^+/-^ mice were maintained on an OF1 genetic background as described in Sonnier et al., 2007 [8–11].

### Purification of recombinant alphα-synuclein and assembly of pre-formed fibrils

Recombinant human α-syn was purified similarly to a previously published protocol [15, 16]. Human α-syn was used in this double-hit model with to provide a higher-fidelity platform for potential downstream pre-clinical screening of α-syn targeting therapeutics. Briefly, the protein was expressed in BL21 *E.coli* transfected with a pT7-7 expression plasmid expressing human α-syn. Once expressed, cells were collected and stored at −80°C. For purification, cells were lysed by sonication and boiling, and centrifuged to remove cell debris. The α-syn-containing supernatant was dialyzed overnight in 10 mM Tris, pH 7.5, 50 mM NaCl, and 1 mM EDTA, using SnakeSkin Dialysis Tubing MWCO 7,000 (Thermo Scientific). Chromatographic separation was performed using a Superdex 200 Column (GE Healthcare Life Sciences) and a Hi-trap Q HP anion exchange column (GE Healthcare Life Sciences). Fractions containing α-syn were identified by SDS-PAGE and Coomassie staining, and then dialyzed overnight into PBS buffer (Life Sciences). A NanoDrop 2000 (Thermofisher) was used to determine the protein concentration by OD280 reading and protein was concentrated to 5 mg/mL using a Vivaspin protein concentrator spin column with a MWCO of 5kDa (GE Healthcare). Aliquots of 500 μL were stored at - 80°C until use. For amyloid fibril assembly, purified recombinant α-syn was thawed and subjected to continuous shaking at 1,000 r.p.m at 37°C in a Thermomixer (Eppendorf) for 7 days. Fibril morphology was confirmed by imaging using a Tecnai G2 Spirit TWIN transmission electron microscope (FEI Company) at 120kV and Thioflavin T (Sigma) fluorescence reading (data not shown). Fibrils were aliquoted and frozen at −80°C until use.

### Stereotactic injections

Prior to injection, human α-syn PFFs were thawed and then sonicated at RT with a water-bath cup-horn sonicator (Misonix XL2020, 50% power, 120 pulses 1 s ON, 1 s OFF). Post-sonication fibril morphology was validated by TEM imaging protocol as described above. Prior to injection, 4-week-old mice (both sexes) were anesthetized with an isoflurane/oxygen mixture and injected unilaterally with either 2 μL of PFFs (5 μg/μl) or 2 μL of PBS as a control in the dorsal striatum (coordinates from bregma: AP: + 0.2 mm; ML: −2.0 mm; DV: - 2.6 mm from dura) at a rate of 0.2 μL/min using a glass capillary attached to a 10 μL Hamilton syringe. After injection, the capillary was left in place for 4 minutes before being slowly removed.

### Euthanasia

At 8 weeks post PFF-injection, mice were deeply anesthetized with sodium pentobarbital and perfused transcardially with 0.9% saline followed by 4% paraformaldehyde in phosphate buffer. Brains were collected and post-fixed for 24 hours in 4% PFA, and then stored at 4°C in 30% sucrose in phosphate buffer until sectioning.

### Histology

Brains were then sectioned into 40 μm-thick free-floating coronal sections using a freezing microtome. Brain sections were stored in cryoprotectant and quenched with sodium periodate. During the staining protocols, sections were incubated with primary antibodies for tyrosine hydroxylase (mouse anti-TH, ImmunoStar) at 1:1000 dilution and p-S129 α-syn (rabbit anti-pS129, Abcam [EP1536Y]) at 1:1000 dilution. Sections were incubated with respective anti-mouse or anti-rabbit biotinylated secondary antibodies (Vector Laboratories) and conjugated with an ABC-HRP biotin/avidin complex kit (Vector Laboratories). Staining was developed with 3,3′-diaminobenzidine and sections were mounted for imaging and analysis. Representative images of pS129 pathology were acquired at 20x magnification using a NIKON Eclipse Ni-U microscope.

### *pS129* α-synuclein scoring

The α-syn inclusions (cell body and neurites) were detected following pS129 staining of a whole series of coronal sections (at 240 μm intervals) per animal. The stained slides were blind-coded prior to analysis. We evaluated pS129 inclusions in 8–10 animals per experimental group. Using the 20x magnification on a NIKON Eclipse Ni-U microscope, we evaluated all sections individually and assigned scores from 0 to 4 based on the relative levels of pS129-positive inclusions (such that 0 = no aggregates, 1 = sparse, 2 = mild, 3 = dense, 4 = very dense), as previously described [17, 18]. The pathology depicted in the brain regions in Fig. 5A are representative of the average score of animals in each experimental group. We also used the average score values in each brain region per group to generate a heatmap (Fig. 5B) to represent the entire dataset.

### Stereology and Densitometry

To evaluate the statistical significance of the observed α-syn pathology, we counted α-syn inclusions with unbiased stereological counting methods using the optical fractionator probe in StereoInvestigator (v10.40, MBF Biosciences). Here, full series of brain sections (5-7 sections) spaced at 240 μm were analyzed per animal and a total of five to seven animals per group (all n ≥ 5 per group) were included. To count the pS129+ labeled cell bodies, contours of each region of interest (SN, VTA, BLA and motor cortex) were drawn using a 4x objective. In the VTA, quantifications were accomplished with 60x magnification using a counting frame of 90 μm X 90 μm and grid size of 180 μm (medio-lateral) and 180 μm (dorso-ventral), along with a guard zone of 2 μm and a dissector height of 15 μm. For each region of interest, these quantification parameters were adjusted to assess at least 300 total cells per animal and per side, with the error of coefficient (m = 1) being less than 0.1. Stereology on TH+ neurons in the SN was performed on sections spaced 480 μm with a 70 μm X 70 μm counting frame, a grid size of 221 μm X 221 μm, guard zone of 1 μm, and dissector height of 15 μm. Striatal TH+ optical density was measured using Scion Image (v.163, Meyer Instruments) and imaged on an Olympus BH-2 microscope (Olympus). Densitometric measurements were averaged from a minimum of 2 sections spaced 480 μm apart.

### Statistics

All graphical data are presented as mean ± SEM. Stereological count data were analyzed using negative binomial mixed-effects regression with zero-inflated adjustments in regions with negligible count data. Densitometric measurements were assessed using a linear mixed-effects regression with a random intercept for each animal subject. All statistical analyses were performed using GraphPad Prism (v7.03) or R (v3.4.4). Significant findings and specific statistical analyses are listed in the figure legends.

## Results

### Study design

The study was performed utilizing the *En1*^+/-^ (OF1 background) mouse line that features a knock-in LacZ cassette in one allele of *En1* with WT levels of Engrailed 2 (Fig. 1A) [9]. The most recent characterization of the *En1*^+/-^ mouse model [7, 8, 19] revealed multiple indices of pathophysiology that follow clinical features of PD. Critically, *En1*^+/-^ animals exhibit downregulation of mitochondrial complex I proteins, progressive loss of midbrain dopamine neurons (more pronounced in A9 than A10 neurons), dysregulation of autophagic markers, axonal pathology and neuroinflammation (Fig. 1B) [7]. We tested the hypothesis that the aforementioned nigrostriatal pathology in the *En1*^+/-^ mouse would lead to an exacerbation of α-syn aggregate formation in mice where the synucleinopathy is triggered by injection of α-syn fibrils. Thus, we generated PFFs according to established protocols, and verified them by transmission electron microscopy (Fig. 1C). To induce α-syn-mediated proteinopathy, we injected 4-week old En1^+/-^ mice with PFFs unilaterally into the striatum. After 2 months we euthanized the mice and conducted neuropathological examinations of their brains using markers for α-syn pathology and dopamine neuron survival (Fig. 1D, E).

**Fig.1.**
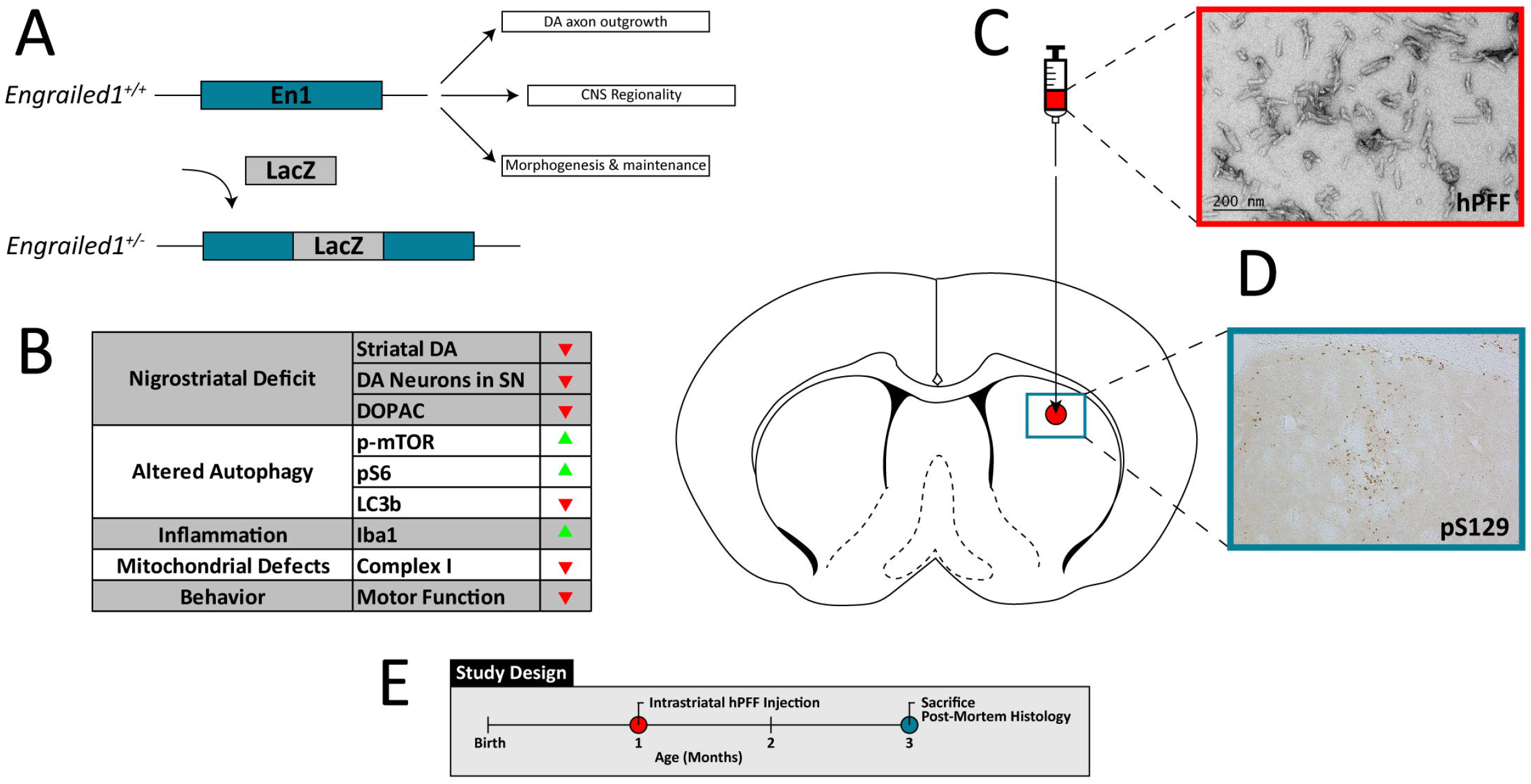
Inducing α-syn pathology in *En1*^+/-^ mice. Engrailed1 expression is vital for CNS development and maintenance, particularly for nigrostriatal projections. **A)** *En1*^+/-^ mice feature a LacZ insertion cassette in one allele and feature **B)** progressive “dying-back” nigrostriatal degeneration, dysfunctional mechanisms in the autophagy-lysosomal pathway, and pathologic neuroinflammatory activation. *En1*^+/-^ mice also feature mitochondrial respiratory chain perturbations and motor dysfunction associated with PD clinical pathology (Nordström *et. al*., 2015; Sonnier *et. al*., 2007) [7, 9]. **C)** Human pre-formed α-syn fibrils were sonicated, validated by TEM, and injected into the dorsal striatum. **D)** PFF injection causes robust pS129 deposition at the site of injection. **E)** Both WT and *En1*^+/-^ animals were unilaterally injected at one month of age and sacrificed for post-mortem histology eight weeks following PFF injection.

### α-Synuclein pathology is exacerbated following injections of fibrils in En1^+/-^ mice

At eight weeks after unilateral, intrastriatal injection of PFFs into *En1*^+/-^ and WT mice, we assessed the numbers of α-syn-positive puncta in four regions with direct projections to the striatum – the SN, BLA, motor cortex and VTA (Fig. 2, 3). We used unbiased stereology to count cell bodies immunolabeled with p-S129-α-syn. The *En1*^+/-^ mice injected with PFFs (En1-PFF mice) displayed a greater than three-fold increase in the number of p-S129-α-syn-positive inclusions in the SN, ipsilateral to the injection of PFFs, compared to WT mice injected with an identical amount of PFFs (WT-PFF mice) (1243 ± 262 cells and 406 ± 52 cells, respectively, p=0.009) (Fig. 2A, B). The numbers of p-S129-α-syn-positive inclusions in the contralateral SN were negligible (Fig. 2B). We observed no Lewy-like α-syn pathology in any PBS injected control animals. To assess the effects of α-syn pathology on nigrostriatal integrity, we assessed the number of dopaminergic neurons in the SN (TH+) and the optical density of TH+ fibers in the striatum. We observed a significant main effect in the loss of striatal fiber innervation caused by the En1 genotype (p=0.001) as expected, but did not observe any added significant effect due to injection of PFFs (Fig. 2D). At the level of the substantia nigra, however, the En1^+/-^ animals (regardless of whether they were injected with PBS or PFFs) did not exhibit significant loss of nigral TH-immunoreactive neurons (Fig. 2C).

**Fig.2.**
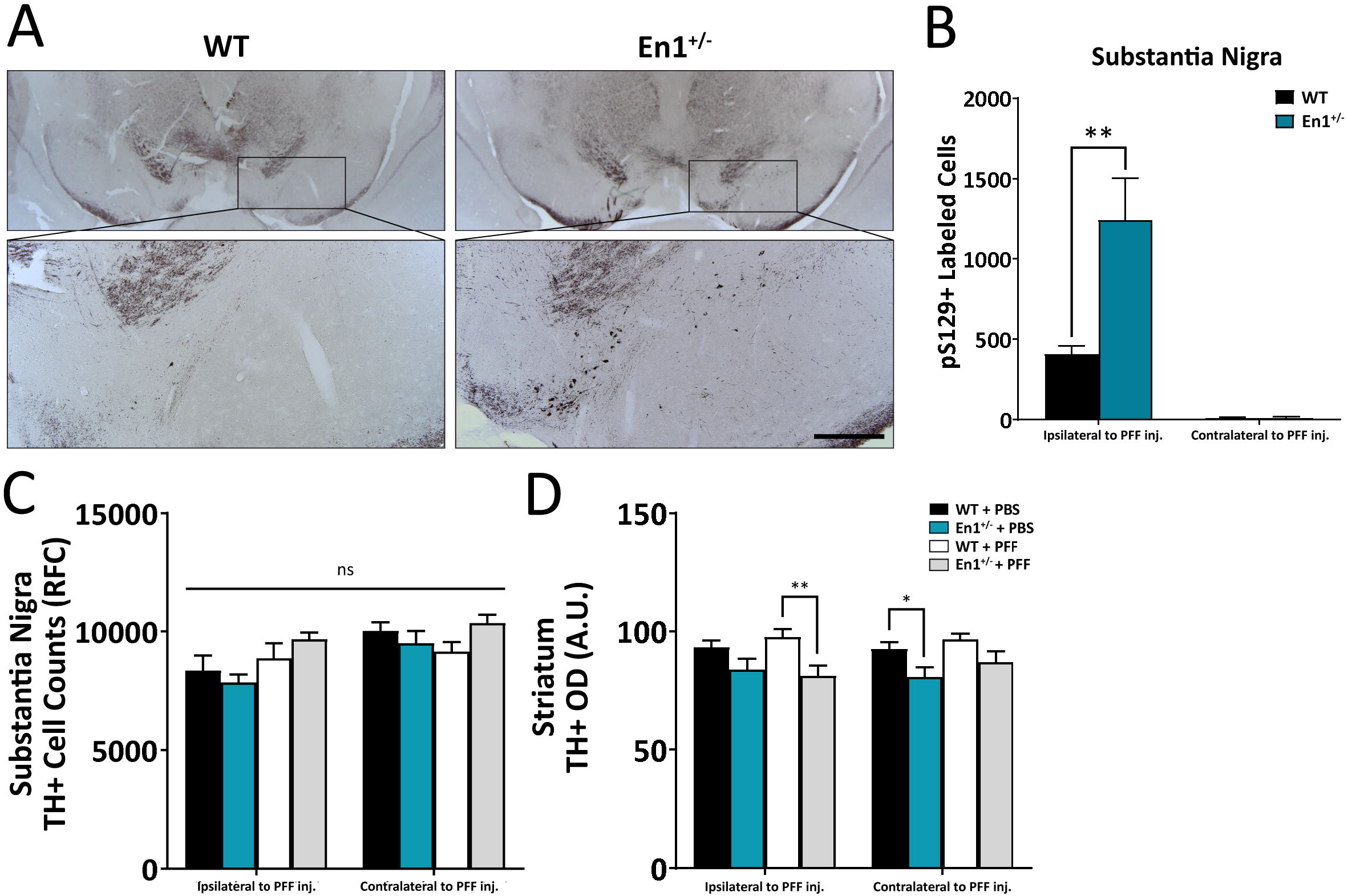
α-Syn pathology is preferentially exacerbated along the nigrostriatal tract. **A)** Microphotographs of α-syn pathology (pS129-immunoreactive cell bodies and neurites) in the substantia nigra of WT and *En1*^+/-^ animals unilaterally injected with PFFs at one month of age. All microphotograph data are representative images from one animal from each respective cohort. **B)** Stereological estimates of pS129+-labeled cells in the substantia nigra ipsilateral and contralateral to the site of PFF injection (N≥8). No α-syn pathology across any stereologically evaluated regions was observed in PBS-treated animals. **C)** Stereological estimates of TH+ labeled cells in the substantia nigra in both WT and En1^+/-^ mice injected unilaterally with either PBS or PFFs. **D)** Corresponding densitometric quantification of TH-immunoreactive striatal integration (N≥5). Results are depicted as mean±SEM. pS129+ and TH+ cell count data were analyzed by negative binomial mixed-effects regression with separate analyses performed to compare pathology ipsilateral to injection and contralateral to injection. Striatal TH^+^ optical density data were analyzed by linear mixed-effects regression. Statistics: all \**p*<0.05, \*\**p*<0.01. Scale bar: A - 250μm.

In the ipsilateral BLA, there was no significant difference in α-syn pathology between *En1*-PFF and WT-PFF mice (2202 ± 476 cells and 1046 ± 208 cells, respectively, p = 0.428, Fig. 3A, B). We noted that the variance was large, and when we analyzed the impact of biological sex we found a marked sex difference, with male *En1*^+/-^ mice displaying 359% increase (p=0.007, compared to WT-PFF mice) in pS129+ deposition in the BLA ipsilateral to PFF injection and females exhibiting a non-significant trend for a 14% increase (p=0.350) (Fig. 3B). In the VTA, similar to the SN, we observed a greater than 3-fold increase in cells with α-syn pathology from the ipsilateral side of *En1*-PFF mice relative to WT mice (821 ± 218 cells and 264 ± 34 cells, respectively, p = 0.033), with negligible pathology in the contralateral side (Fig. 3C). In the motor cortex, we observed no statistically significant difference in α-syn pathology between *En1*-PFF and WT-PFF mice (Fig. 3D).

**Fig.3.**
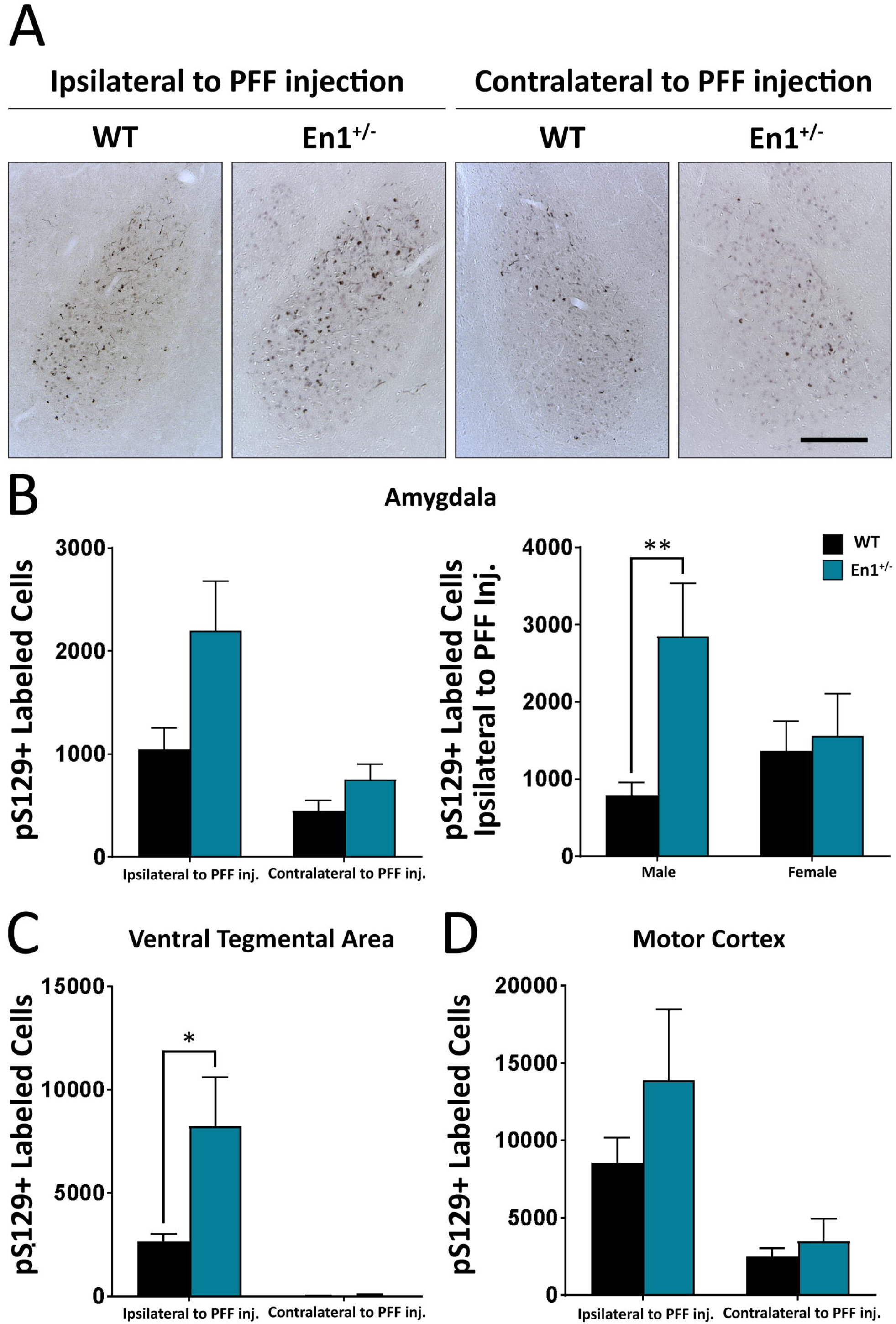
Enhanced propagation of α-Syn pathology observed in extra-nigrostriatal brain regions of *En1*^+/-^ mice. **A)** Microphotographs of α-syn pathology in the basolatereal amygdala of WT and *En1*^+/-^ animals unilaterally injected with PFFs. All microphotograph data are representative images from one animal from each respective male cohort. **B)** Corresponding stereological estimates of pS129^+^ labeled cells in the basolateral amygdala ipsilateral and contralateral to site of PFF injection (N≥8). The second amygdala graph depicts pS129^+^ labeled cells ipsilateral to site of PFF injection separated by sex of animals. **C)** Stereological estimates of pS129^+^ labeled cells in the ventral tegmental area (N≥5) and **D)** motor cortex (N≥6). Results are depicted as mean±SEM. pS129^+^ count data were analyzed by negative binomial mixed-effects regression with separate analyses performed to compare pathology ipsilateral to injection and contralateral to injection. Statistics: all \**p*<0.05, \*\**p*<0.01. Scale bar: A - 250μm.

To map long distance α-syn pathology propagation in the *En1*-PFF mice, we used a semi-quantitative scale and determined a mean pathology score based upon p-S129-α-syn-positive perikarya and neurites in all brain regions where α-syn inclusions were detectable. Enhanced pathology (both ipsilateral and contralateral to PFF injection) was observed in *En1*-PFF mice across a number of cortical regions, as depicted in Figure 4. The pathology scoring data is summarized in brain- and heat-maps (Fig. 5A, B). While there was less pathology in the olfactory bulb, we observed increased α-syn pathology in the frontal association, orbital, motor, ectorhinal, somatosensory, perirhinal, auditory, and entorhinal cortices of both *En1*-PFF and WT-PFF mice (Fig. 4), primarily ipsilateral to the intrastriatal PFF injection, with more pronounced pathology observed in *En1*-PFF mice (Fig. 4, 5).

**Fig.4.**
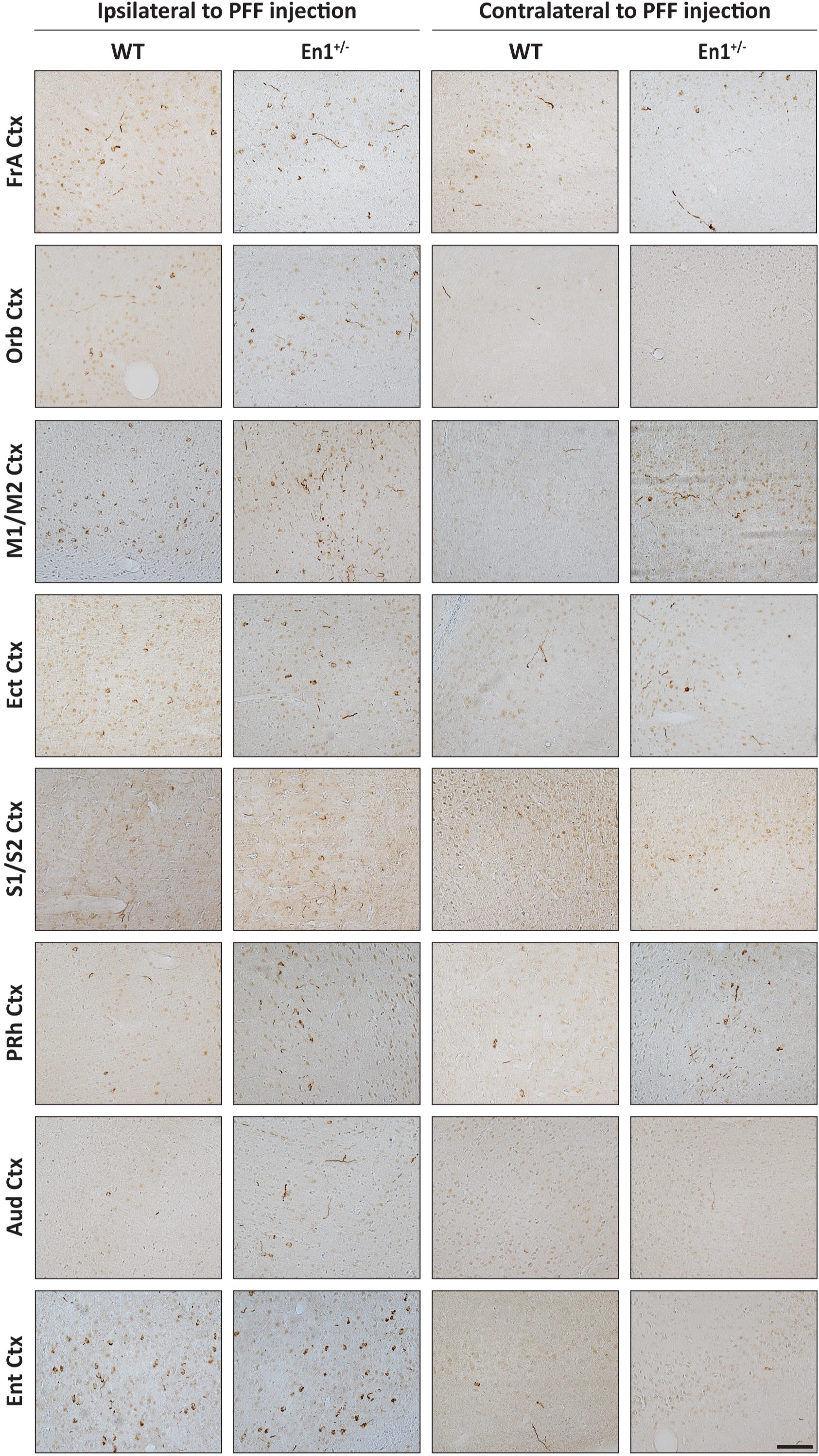
Striatal PFF injections induce widespread cortical α-Syn pathology in *En1*^+/-^ mice. Representative microphotographs of pS129 staining across cortical brain regions in both WT and *En1*^+/-^ mice. α-Syn pathology in cell bodies and neurites is detected across cortical regions with both direct and indirect projections to/from the striatum. Pathology is observed both ipsilateral and contralateral to PFF injection. Data are representative images from one respective animal of each cohort. Abbreviations: FrA – Frontal Association, Orb – Orbital, M1/M2 – Motor, Ect – Ectorhinal, S1/S2 – Somatosensory, PRh – Perirhinal, Aud – Auditory, Ent – Entorhinal. Scale bar: 500μm.

**Fig.5.**
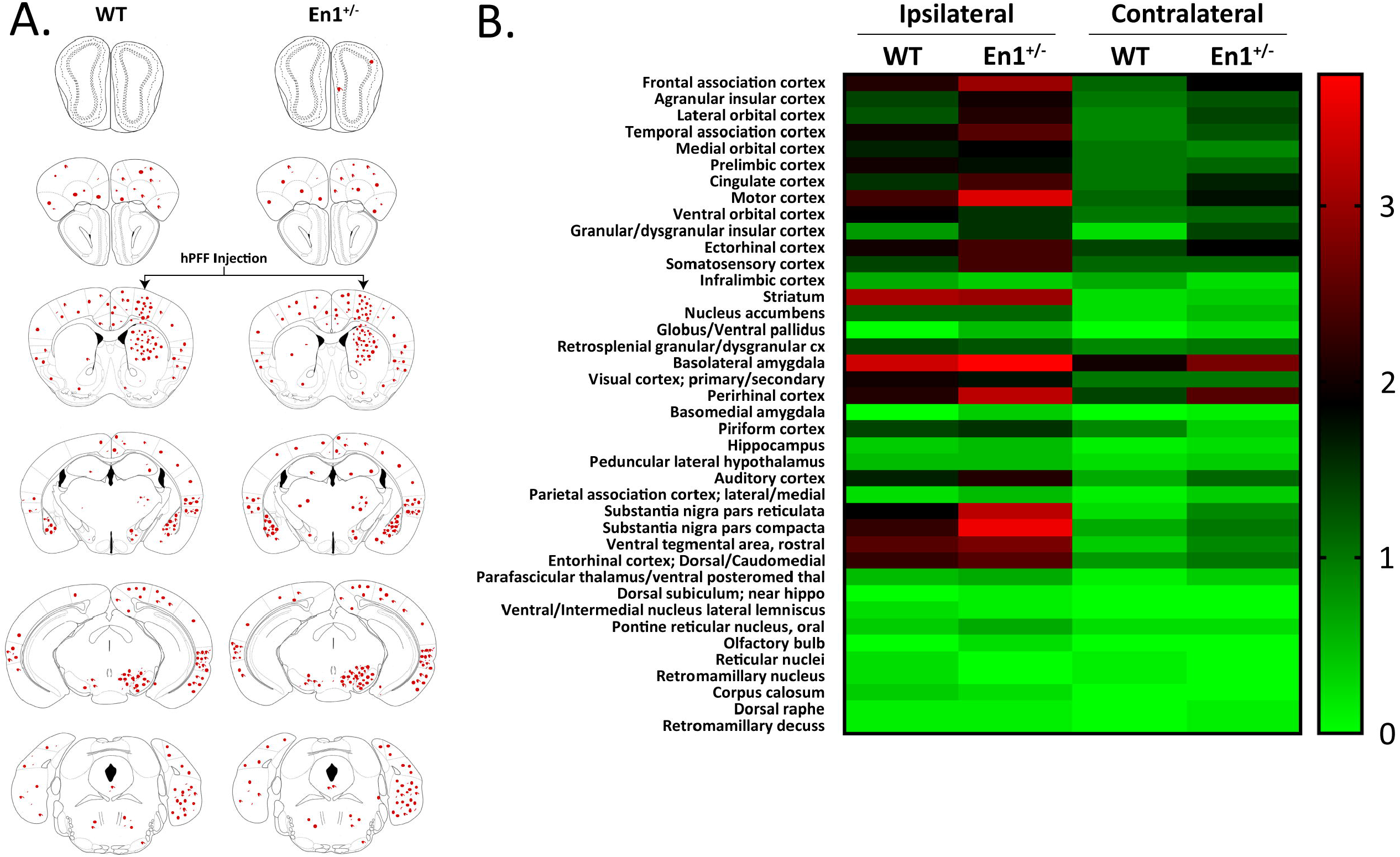
Pathological propagation of α-Syn is enhanced throughout *En1*^+/-^ neuraxis. **A)** Schematic representation of brain regions depicting sites of observed pS129 cellular and neuritic deposition. **B)** Heat map representing semi-quantititative pS129-pathological scores across numerous regions in both WT and *En1*^+/-^ mice injected with striatal PFFs (N=8). Regions were assessed both ipsilateral and contralateral to PFF injection.

## Discussion

We quantified p-S129-α-syn-positive pathology following injection of α-syn PFFs into the striatum of mice and found that the absence of one *En1* allele led to significantly increased numbers of cells displaying α-syn aggregates in the SN and VTA, as well as in the BLA of male mice. Qualitatively, we also observed more p-S129-α-syn-positive staining in several cortical regions that project to the striatum. We have previously investigated many facets of neuropathology in *En1*^+/-^ mice and showed that this mouse model exhibits several neuropathological features resembling those seen in PD, but lacks α-syn aggregates [7]. Unusual features of the *En1*^+/-^ mouse that are particularly pertinent to PD are the protracted degeneration of nigral dopamine neurons over several months, with a later and less pronounced loss of dopamine neurons in the VTA, as well as the early onset reductions in evoked striatal dopamine release and a conspicuous axonal pathology reminiscent of the axonal dying back that is believed to occur in PD [7, 20]. Notably, the first signs of axonopathy precede nigral degeneration in *En1*^+/-^ mice, adding to their relevance to PD where loss of dopaminergic terminals in the putamen also has been shown to precede nigral cell loss [21].

There are many potential factors that could explain how heterozygous loss of *En1* exacerbates pS129-α-syn pathology. Mitochondrial deficits in midbrain dopamine neurons that occur in *En1*^+/-^ mice may aggravate fibril recruitment of endogenous α-syn, and subsequent aggregation and propagation. Deficiencies in mitochondrial complex I proteins, Ndufs 1 and 3, as well as critical mitochondrial homeostatic mediator, LaminB2, are signature hallmarks in *En1* model pathophysiology [8, 22]. *En1*^+/-^ mice also feature robust oxidative stress, most likely attributed to signaling downstream of mitochondrial impairment or to increases in LINE-1 expression and DNA damage [11, 12, 23]. Oxidative damage in En1-expressing neurons may induce enhanced α-syn modification and accelerate proteopathic seeding capacity.

Although the increase in p-S129-α-syn labeling was most prominent in the SN of the *En1*^+/-^ mice, a significant increase in p-S129-α-syn-positive inclusions was also apparent in the ipsilateral VTA. It is possible that some of the injected PFF, which was targeted into the central striatum, may have diffused into the ventral striatum leading to uptake of α-syn fibrils by terminals from dopaminergic neurons located in the VTA. The deposition of p-S129-α-syn was not as pronounced in VTA neurons as in the SN, possibly due to the anatomical coordinates of PFF injection into the striatum primarily leading to targeting of central and dorsal striatal regions innervated by substantia nigra and not VTA. It is also possible, however, that dopaminergic neurons in the SN are more impaired due to reduced *En1* levels than those in the adjacent VTA dopamine neurons, which would be consistent with earlier reports of greater dopamine neuron loss in the nigra compared to VTA in *En1*^+/-^ mice [9, 24]. We did not observe reduction in the number of TH-immunoreactive neurons in the SN of *En1*^+/-^ mice; however, we detected significant decreases in striatal fiber innervation which are known to precede SN neuron loss both in the En1 heterozygous mouse and in clinical PD [7, 21]. Although SN neuron loss has been previously reported at three months of age [9], genetic background is known to significantly impact the extent of degeneration of nigral dopamine neurons [20], and subtle differences in such genetic susceptibility loci between our present *En1*^+/-^ colony and those used in earlier papers could potentially have affected the results. Notably, the intrastriatal injections of PFFs did not cause any additional striatal degeneration or accelerated nigral neuron loss. This is not surprising considering that our follow up after PFF injection was only 2 months (mice aged 3 months at the time of sacrifice) and earlier work indicates that in WT mice the loss of nigral TH-positive neurons does not become significant until between 3 and 6 months after intrastriatal PFF injection, reaching 35% at 6 months [15]. We opted to inject the PFFs into *En1*^+/-^ mice that were only one-month old, preceding the time when they exhibit extensive nigrostriatal axonal degeneration, to ensure that the nigrostriatal neurons were still capable of uptake and transport of the PFFs. Future studies with longer survival times are necessary to determine whether the PFF injections cause nigral neuron death that is additive to the nigral cell death that occurs in *En1*^+/-^ mice (estimated to be around 40% at 6 months of age), or if it is the identically vulnerable subpopulation of nigral dopamine neurons that dies in both paradigms.

Interestingly, we also observed increased α-syn deposition in the BLA of male En1^+/-^mice. It is unlikely that the increased α-syn pathology in the BLA is primarily due to cell-autonomous deficits in BLA neurons because there is no reported evidence of *En1* expression in the adult amygdala. Notably, neurons in amygdala project to both SN and striatum [25]. Thus, the increased α-syn pathology could be due to mitochondrial deficits in nigrostriatal dopamine neurons in *En1*^+/-^ mice resulting in a greater amount of aggregated α-syn seeds, or the neuroinflammatory environment induced by neurodegeneration in the SN and striatum. While we observed a significant exacerbation of pathology in the BLA of male *En1*^+/-^ mice injected with PFFs, compared to WT mice similarly injected with PFFs, it is unclear why female *En1*^+/-^ mice did not show a similar increase in proteinopathy compared to WT female mice. There are no previously reported sex differences in phenotypes of *En1*^+/-^ mice, and it is unclear that the impact of biological sex on the neuropathology of *En1*^+/-^ mice has ever been addressed. It is likely that due to the relatively short delay between PFF injection and analysis of neuropathology in our study, female mice may develop equivalent levels of amygdala pathology latently. The divergence in pathology may also be a result of intrinsic sex differences in amygdala networks and gene expression, previously described in rodents, that may alter vulnerability to proteinopathy [26, 27]. Future studies will require further investigation of sexual dimorphism pertaining to the *En1* allele loss and how it may affect relevant brain structures.

Using stereological analysis, we found a non-significant trend for an increase in p-S129-α-syn-positive aggregates in the ipsilateral motor cortex of the *En1*^+/-^ mice. Bilaterally, in several other neocortical brain regions which also project to the striatum and where we did not conduct formal quantification of α-syn pathology, the overall visual impression was that there was more p-S129-α-syn staining in the *En1*^+/-^ mice (Fig. 4, 5 A, B). While we did not attempt to quantify pathology in all the cortical regions and cannot judge whether there were robust effects of the absence of one *En1* allele, it is possible our study was underpowered to detect differences in α-syn pathology between the *En1*^+/-^ and WT mice in the motor cortex. It is also likely that the precise location of the PFF injections in the striatum, which receives topographically region-specific corticostriatal projections, contributed to a high variability that masked a potential effect.

Taken together, we suggest that the *En1*-PFF mouse represents a potentially powerful PD mouse model, with many molecular and neuropathological features resembling PD. Previous descriptions of the *En1*^+/-^ mouse have reported that it exhibits progressive, retrograde degeneration of dopamine neurons focused in the SN and less marked in the VTA [7, 9] as well as reduced striatal levels of dopamine, neuroinflammation, and changes in markers of autophagy [7]. In the current study, we found that the additional hit of a unilateral intrastriatal PFF injection provides α-syn aggregate pathology which is pertinent to PD pathology, and which was enhanced in *En1*^+/-^ mice compared to wild type mice. The combination of all these findings makes the *En1*-PFF mouse relevant for use in the testing of experimental therapeutics. Further studies are needed with bilateral intrastriatal injections of PFFs and long-term survival of animals to establish how severe the nigral degeneration and synucleinopathy can become, and whether the mice develop a range of progressive motor and non-motor deficits that mimic those seen in PD develop.

## Acknowledgments

We acknowledge the Van Andel Research Institute and the many individuals and corporations that financially support research into neurodegenerative disease at the Institute. D.S.S. was supported by a José Castillejo Mobility Grant (CAS15/00201) awarded by the Ministry of Education, Culture and Sport (Madrid, Spain). N.L.R.’s work is supported by the Peter C. and Emajean Cook Foundation. J.M. is supported by National Institutes of Health grants (R21NS101676 and R01NS060729). J.H.K. is supported by a Rush University center grant from the Parkinson’s Disease Foundation. J.H.K. reports additional support from the National Institutes of Health, the Michael J. Fox Foundation, and the Multiple System Atrophy Coalition which are outside but relevant to the submitted work. P.B. is supported by grants from the National Institutes of Health (1R01DC016519-01 and 5R21NS093993-02). J.M., P.B. and J.H.K. are supported by an NIH IGNITE grant (1R21NS106078-01A1). P.B. reports additional grants from Office of the Assistant Secretary of Defense for Health Affairs (Parkinson’s Research Program, Award No. W81XWH-17-1-0534), The Michael J Fox Foundation, National Institutes of Health, Cure Parkinson’s Trust, which are outside but relevant to the submitted work. We additionally acknowledge the technical expertise of Lindsay Meyerdirk and Emily Schulz in their contributions to this study.

